# STRIDER: <u>St</u>e<u>ri</u>c hin<u>d</u>rance <u>e</u>stimato<u>r</u>

**DOI:** 10.1101/2020.02.07.931550

**Authors:** L Ponoop Prasad Patro, Thenmalarchelvi Rathinavelan

## Abstract

*In silico* modeling plays a vital role in the *de novo* designing and docking of biomacromolecules as well as in exploring their conformational dynamics. Additionally, it has a major role in acquiring the structural insights from the parameters derived from the experimental techniques such as cryo-electron microscopy. Steric hindrance is one of the important measures to validate the accuracy of the constructed model. A web user interface (WUI) namely, STRIDER (<u>st</u>e<u>r</u>ic h<u>i</u>n<u>d</u>rance <u>e</u>stimato<u>r</u>) (www.iith.ac.in/strider/) can estimate and report pairwise inter- and intra- molecular steric hindrances using the van der Waals radius of 117 elements through a user interactive interface. STRIDER also identifies and reports the coordination number of 64 metals along with their interacting pattern in an interactive mode. STRIDER can analyze an ensemble of conformers, wherein, multiple conformers are used to circumvent sampling issue in flexible docking, understand protein folding and facilitate structure based virtual screening. Further, it generates a pymol session file that can be used for offline analysis. As STRIDER simply requires the Cartesian coordinates of the given molecule in protein data bank format, any chemical structure can be an input.

**Availability:** It can be freely accessible through: www.iith.ac.in/strider/ without any registration.

**Theme Of the Concept:** 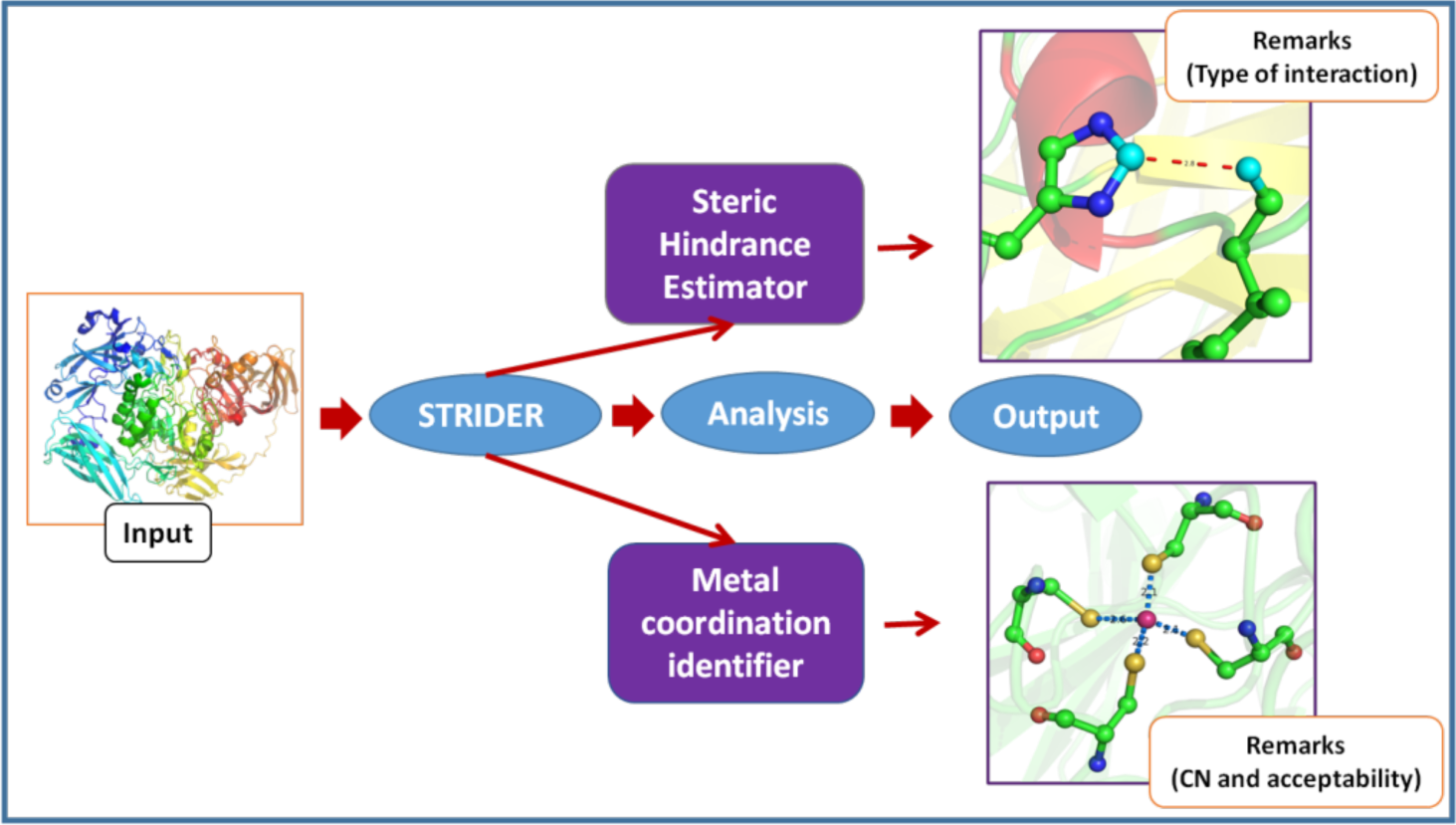

## Introduction

Biomacromolecules like carbohydrates, nucleic acids, proteins and lipids are the pivotal building blocks of the cell structure and play key roles in the organelle & cellular functions and inter-organelle & inter-cellular communications. Establishment of their roles in the formation & function of organelles and cells under normal and diseased states necessitate atomistic details about the biomolecular structure and dynamics. As there are limitations in exploring these biomolecular structures through experimental techniques like X-ray crystallography, NMR spectroscopy and cryo-electron microscopy, molecular modeling becomes a promising strategy. For instance, molecular modeling acts as a supplement to experimental techniques to derive structural and functional information of supramolecular nanomachines like bacterial secretion systems(Rathinavelan et al., 2010, Dey et al., 2019) and bacterial surface associated polysaccharide exportation machineries(Sachdeva et al., 2016, Bushell et al., 2010, Hsu et al., 2017, Patel et al., 2017, Pavlova et al., 2016, Sachdeva et al., 2017, Patro and Rathinavelan, 2019) to establish the associated functional mechanisms involved in virulence exportation and host…pathogen interactions. Further, the inter-molecular nucleic acids triplexes are not generally amenable to structure determination by experimental techniques although they have been presumed to have a vital function in gene regulation besides their promising role in antigene therapy(Moser and Dervan, 1987),(Le Doan et al., 1987). Thus, molecular modeling is crucial in establishing the biological significance of nucleic acids triplexes(Thenmalarchelvi and Yathindra, 2005, Rathinavelan and Yathindra, 2006). Similarly, to establish (i) the rationale behind the inefficiency of mismatch repair proteins to act on noncanonical base pair mismatches(Khan et al., 2015, Rossetti et al., 2015) under disease conditions, (ii) the sequence dependent influence of various base pair step and base pair parameters in chromatin folding(Tolstorukov et al., 2007) and (iii) the role of oligosaccharides in host…pathogen interaction(Varki, 1993), molecular modeling is essential.

All of the above require a good starting model with accurate internal geometries, which is also free from steric hindrance (synonymously, short contact, bad contact or bump) and lies close to the local minima in the energy landscape. To this end, we have developed a web server namely, STRIDER (<u>st</u>e<u>r</u>ic h<u>i</u>n<u>d</u>rance <u>e</u>stimato<u>r</u>) (www.iith.ac.in/strider/) that calculates intra- & inter-molecular interatomic distances and reports the steric hindrance, if at all any. As the RNA structure prediction(Flores et al., 2010), homology modeling(Ashkenazy et al., 2011, Kundrotas et al., 2008), flexible docking(Meng et al., 2011, Sliwoski et al., 2014), structure based virtual screening(Yan et al., 2017), multiple conformer generation of the lead molecules in cheminformatics(Hawkins, 2017), modeling intermediate conformers to understand protein folding(Burnley et al., 2012, Kingsley and Lill, 2014, Jamroz et al., 2016) *etc.* may require multiple conformers to overcome the sampling issues, STRIDER accepts ensemble of conformers and reports pairwise steric hindrance profile for individual conformers in an interactive manner. Although the metal ion coordination has an important role in biomolecular stability(Largy et al., 2016, Zhang et al., 2014, Bhattacharyya et al., 2016) and function(Riordan, 1977, Pattammattel et al., 2013, Liu et al., 2014), identifying metal ions directly by NMR and X-ray crystallography still remains a challenge(Jensen et al., 2005, Erat and Sigel, 2011, Nabuurs et al., 2006, Handing et al., 2018), thus, calls for metal modeling. Thus, when a query structure with metal ion is submitted to STRIDER to access the accuracy of the modeled metal coordination, it reports the metal coordination number and distance profile of metal with its coordinating partners in the binding pocket. Many online and offline tools such as Cytoscape(Shannon et al., 2003), RING2.0(Piovesan et al., 2016), RINalyzer(Doncheva et al., 2011), PLIP(Salentin et al., 2015), Arpeggio(Jubb et al., 2017), Ligdig(Fuller et al., 2015), GIANT(Kasahara and Kinoshita, 2014), PyMOL (www.pymol.org), Joy(Mizuguchi et al., 1998), Bioptools(Porter and Martin, 2015), PIC(Tina et al., 2007), CheckMyMetal(Zheng et al., 2017) and LIGPLOT+(Laskowski and Swindells, 2011) are available to characterize favorable and unfavorable interactions of biomolecules, but, each with their own advantages and disadvantages. STRIDER can be a good addition to these resources by providing pairwise unfavorable contacts and metal interaction profiles for a range of conformers in a user-friendly interactive manner.

### STRIDER architecture

The user interface of STRIDER server is developed using PHP web server scripting language, HTML and Java script. It has two modules for structural analyzes of the biomolecules: (i) steric hindrance estimator and (ii) metal binding identifier. The methodologies used by these applications are discussed below. The input for the STRIDER server is a three-dimensional coordinate of the molecule of interest in the protein databank (PDB) format and output is provided in the result page as described below.

### Steric hindrance estimator

The steric hindrance estimator executes a python based program to create a primary profile of the intra- and inter-molecular atom pairs that fall within 5Å distance in the query molecule. The created pairwise primary profile is used in the subsequent distance calculation, wherein; the calculated distance is compared with the sum of van der Waals radii of the corresponding atoms. For the creation of pairwise distance profile of each ‘i^th^’ atom, i+3 to n^th^ (wherein, ‘n’ is the total number of atoms in the system) atoms are considered in incremental direction. Using the primary profile, a secondary profile is created which consists of atom pairs list that fall below 0.4Å lesser than the sum of van der Waals radii (Mantina et al., 2009, Hu et al., 2009). Subsequently, the secondary profile is used to distinguish the steric hindrance from the non-covalent interactions (such as hydrogen bond and hydrophobic interaction) and metal interactions by considering the nature of the atoms (*viz.*, polar, non-polar, metal and electro-negativity)(Chen et al., 2010). For instance, if the distance between a pair of electronegative atoms falls in the range of 2.2-3.5Å, it is categorized under “Hydrogen Bond” (HB). Similarly, if any electronegative atom falls within 3.5Å distance from a metal ion (64 metals), it is considered under “Metal Interaction” (MI) category. Further, the proximity between any two carbon atoms is in the range of 3.0-8.0Å, then it is considered under “Hydrophobic Interaction” (HP) category(Rajgaria et al., 2009). When the atom pairs in the secondary profile do not fall under any of the aforementioned categories, it is classified as “Steric Hindrance” (SH) category.

After the completion of the analysis, STRIDER automatically directs the user to a result page that contains a summary table (**Figure 1**), which reports the number of occurrences of the aforementioned interactions in the query molecule. For the detailed inspection of a specific category, the user has to simply click the corresponding cell in the summary table that is hyperlinked to a second result page, which contains information of all the atom pairs fall under the particular category. A table in the second result page provides elaborated information such as chain name(s), atoms name(s), atom number(s), residue name(s) and residue number(s) that correspond to each atom pairs in a rowwise manner (**Figure 1**). This can be downloaded as a text file and can be utilized for future analyzes. If the user wants to visualize an individual short contact (specific to each atom pair), the hyperlinked cell in the table (the third column of the table) has to be clicked to zoom-in the region of interaction in the JSmol interactive viewer (http://www.jmol.org/), wherein, the distance between the atoms is marked in a dotted line. As each query job is assigned an identification number (IDN), the user can access the data for a week by using the corresponding job IDN.

**Figure 1.**
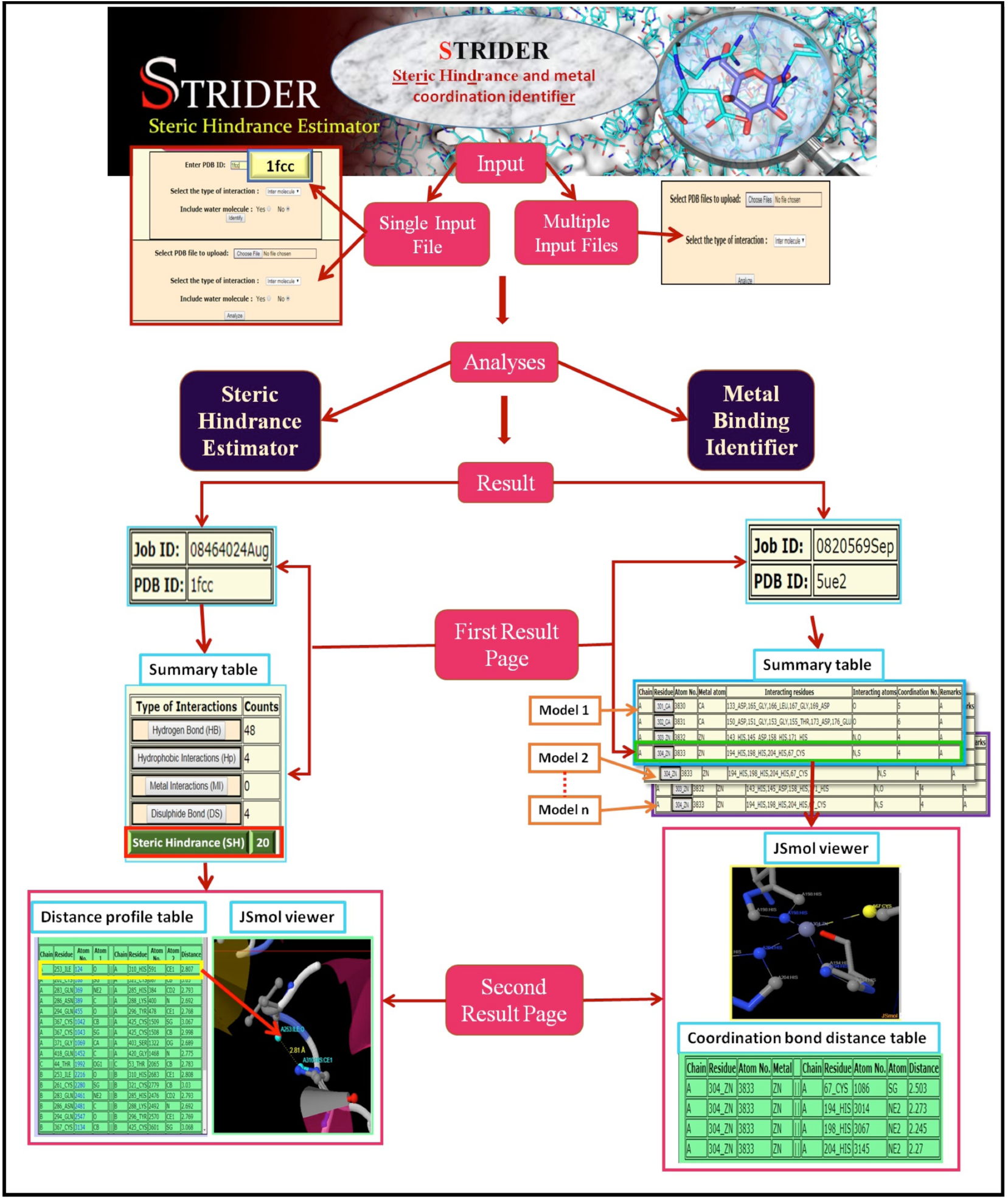
Schematic diagram describing the workflow of “Steric Hindrance Estimator” (Left) and “Metal Binding Identifier” (Right) modules of the STRIDER WUI. (**Left**) In the main result page, different types of interactions present in the PDB ID: 1FCC (query) are given in a summary table. Distance profile corresponding to “SH” (Steric Hindrance) (red colored box) category is displayed in the secondary result page alongside with the zoomed-in view of a selected atom pair (yellow colored box). (**Right**) The metal coordination corresponding to 16 NMR conformations of Ca^2+^ and Zn^2+^ binding matrix metalloproteinase-7 zymogen (PDB ID: 5UE2) is summarized in the first result page. In the secondary result page, the coordination bond distance profile corresponding to Zn^2+^ (residue number 304, green colored box) of model number 1 is shown.

STRIDER can also analyze multiple conformers (currently, restricted to 100 structures) that can be uploaded as a single file (like in NMR format with each conformers is identified by the corresponding ‘MODEL number’ and separated by ‘ENDML’ lines) or as multiple files with any name for each conformer. Due to the requirement of extensive computational time, STRIDER does the multimer steric hindrance estimation always in the background and the user can check the job status by using the job IDN. In the case of multiple conformers in a single file, the results are reported modelwise and the user has to simply select the model number of the interested conformer to see the zoomed-in pairwise steric hindrance profile. In the case of multiple input files, clicking the corresponding file name can access modelwise report.

### Metal binding identifier

STRIDER also extends its application to estimate the metal ion coordination in the query structure. For instance, whenever it encounters the distance between a metal ion and any electronegative atom false less than 3.5Å, it creates a distance profile for further inspection of the metal binding site. Subsequently, the number of such instances corresponding to a particular metal ion is counted and updated in the distance profile. The metal coordination output profile is similar to the steric hindrance output profile and can be downloaded in the text format. The output is summarized in a table with one metal per row and the cells that contain the metal residue number are hyperlinked to a secondary result page. The secondary result page contains the information about the atoms that are coordinating with the respective metal ion and the interaction can be closely visualized in a JSmol interactive visual window. In the JSmol applet window, the metal is represented in a sphere and the binding residues are represented in ball and stick with the coordination bond indicated in the dotted lines. The metal coordination can also be analyzed for the multiple conformers as discussed above.

### Specific examples

To perform steric hindrance estimation, the user has to choose ‘Steric Hindrance’ sub-menu bar in the main-menu bar which will direct the user to choose anyone of the following choices: single file or multiple files (**Figure 1**). The first option takes a single file as the input that contains either one conformer or ensemble of conformers in NMR format and the second option takes multiple files as the input (**Figure 1**), wherein, each file contains a single conformation of the query molecule. The PDB IDs can also be given as input in STRIDER for the analyses. In this case, the coordinates corresponding to the PDB IDs are directly taken from the protein databank. The stepwise execution process of steric hindrance estimator is shown **Figure 1 (Left)**. For instance, the intra-molecular distance profile is computed for PDB ID: 1FCC that contains a single conformation of streptococcal protein G and human IGG complex. Upon completion of the calculation, STRIDER directs the user to the output page, wherein; the results can either be accessed through display or be downloaded. The immediate result page contains a summary table, which carries a link to a secondary result page (red colored box in **Figure 1 (Left)**). The secondary result page contains a distance table with one atom pair per row under the selected category (for example, ‘SH’ in **Figure 1 (Left)**). The interactive zoomed-in view of a specific atom pair can be invoked in JSmol by clicking on the atom number of the first atom in the table (yellow colored box in **Figure 1 (Left)**). The metal binding identifier option is illustrated in **Figure 1 (Right)** by considering PDB ID: 5UE2 as a case in point that contains multimers of Ca^2+^ and Zn^2+^ binding matrix metalloproteinase-7 zymogen.

Based on the generated ‘steric hindrance’/’metal coordination’ profile, a pymol session (.pse extension) file is created in the background with the steric hindrance(s)/coordination bond(s) distance marked. In the case of the multiple conformers, a single pymol session file containing the multiple conformers is created. On the other hand, in the case of multiple files, multiple pymol session files are created. This pymol session file(s) can be downloaded along with the text file containing ‘steric hindrance’/’metal coordination’ profile in a compressed form (tar and zipped form) and can be for visualization in the pymol molecular visualization system.

## Conclusions

Manual modeling comes into picture when the experimental techniques fail to provide the structural information of biomacromolecules. During the modeling, one has to be cautious as the modeled structure should not only be good in terms of internal geometries, but also, be steric free. As the sum of van der Waals radii is the best quantifier to access whether a pair of atoms are in unfavorable proximity or not, we have developed a web user interface, namely STRIDER that can estimate and visually display all the inter- and intra-molecular steric hindrance of a query molecule. It further reports the metal coordination number of metals found with the query structure. Both the estimations can also be performed for an ensemble of structures that are either embedded in a single file or provided as multiple input files. Thus, STRIDER acts as an essential tool in the modeling of any 3D structure in PDB format.

## Author contributions

LPPP developed the web server. TR designed and supervised the entire project. LPPP and TR wrote the manuscript.

## Acknowledgements

The work was supported by BIRAC-SRISTI GYTI award (PMU_2017_010), BIRAC-SRISTI GYTI award (2019), Department of Biotechnology, Government of India: IYBA-2012 (D.O.No.BT/06/IYBA/2012), BIO-CaRE (SAN.No.102/IFD/SAN/1811/2013-2014), and R&D (SAN.No.102/IFD/SAN/3426/2013-2014) and Indian Institute of Technology Hyderabad (IITH). LPPP is supported by MHRD fellowship.

